# Predictive factors of response to electrochemotherapy in canine oral malignant melanoma

**DOI:** 10.1101/727164

**Authors:** Matías Nicolás Tellado, Felipe Horacio Maglietti, Sebastián Diego Michinski, Guillermo Ricardo Marshall, Emanuela Signori

**Affiliations:** Laboratorio de Sistemas Complejos, Departamento de Computación, FECyN, Universidad de Buenos Aires, Ciudad de Buenos Aires, C1428EGA, Argentina; Facultad de Ciencias Veterinarias, área de Química Biológica, Universidad de Buenos Aires, Ciudad de Buenos Aires, C1417DSD, Argentina; Instituto Universitario del Hospital Italiano - CONICET, Ciudad de Buenos Aires, C1199ACL, Argentina; Instituto de Física del Plasma, Departamento de Física, FCEyN, Universidad de Buenos Aires - CONICET, Ciudad de Buenos Aires, C1428EGA, Argentina; CNR-Institute of Translational Pharmacology, Roma, 00133, Italy

## Abstract

Electrochemotherapy is a treatment modality which has been increasingly used in veterinary and human medicine for treating cutaneous and subcutaneous tumors. In this prospective work we evaluated the outcome of using electrochemotherapy as a first-line treatment for canine oral melanoma in different stages, with the aim of determining predictive factors of response to the treatment. Mucosal melanoma is the most common cause of oral cancer in dogs. Canine oral malignant melanoma is very similar to human oral melanoma in many aspects, being a very good translational model for studying response to this treatment. Sixty-seven canine patients were treated. Intravenous bleomycin was the preferred drug, and the standard operating procedures for electrochemotherapy were followed. The patients were followed-up for two years. According to WHO criteria, the objective response per stage was: stage I 100%, stage II 89.5%, stage III 57.7% and, stage IV 36.4%. The overall median survival was 7.5 months (2-30 months, mean 9.1 months). Median overall survival of patients in stage I was 16.5 months, in stage II was 9 months, in stage III 7.5 months, and in stage IV 4.5 months. The average number of electrochemotherapy sessions was 1.5 for every stage. The incidence of new metastases among treated patients was 28.4%. Patients in advanced stages, with bone involvement, and caudal location of the tumor had poorer response rates and shorter overall survival times. The treatment greatly improved the quality of life of the patients. Electrochemotherapy is an important technique in the oncological armamentarium against melanoma, and these results can be used to predict human response to this therapy in each stage.

## Introduction

Spontaneous tumors in dogs have many advantages as a human model of disease, they develop in the presence of an intact immune system through the exposure to the same carcinogens[1]. Human and canine malignant melanoma share many histological characteristics such as expression of melanocyte differentiation antigens, melanocyte morphology, patterns of growth necrosis and ulceration. Both have propensity to metastasize to regional lymph nodes and brain as to other organs[2]. They are suitable for preclinical trials, as the tumors evolve naturally and are very heterogeneous, making the results easier to translate into human medicine. Provided the treatment studied renders good results, there is a benefit for patients and owners.

Oral malignant melanoma represents 6% of the neoplasms in canines. It is the most common oral cancer. Overall survival without treatment is around 65 days[3]. The mean age of diagnostic is 11.6 years[4].Oral malignant melanoma in dogs is more aggressive and has a poorer prognosis than melanomas of other locations[5,6]

Most common breeds affected by melanoma are cocker spaniel, German shepherd, pointer, Weimaraner, golden retriever, poodle, chow-chow and boxer. The disease usually is locally infiltrative with metastatic progression in more than 80% of the cases. On one side, when the disease is diagnosed early in its course, or has a rostral location, there is a better outcome. On the other side, poorer prognosis is associated with a late diagnosis, caudal location, presence of satellite lesions, and when dysphagia and dyspnea are present[7].

First line treatment of oral melanoma is surgical excision with clean margins. But, since the majority of tumors invade bony structures, complete surgical resection is often very difficult. When this is the case, or if it has metastasized to lymph nodes, surgery plus radiotherapy is the treatment of choice. Up to 70% remission rate is achieved with radiotherapy alone. However, recurrence of distant metastases can arise 5 to 7 months after treatment. Coadjuvant chemotherapy does not increase response rates or adds survival benefit to the patients[8].

Recently, new immunotherapies have been developed, including a DNA-based vaccine to control or even eradicate the disease[9]. At the moment, these kinds of treatments are only to be used as adjuvant therapies to surgery, radiation or chemotherapy. They provide a significant improvement in overall survival in patients treated in combination with this approach.

Electrochemotherapy (ECT) is a new treatment modality which has been gaining territory in oncology since the Standard Operating Procedures were published in 2006[10]. ECT consists of the application of an electric field to a tumor in order to increase the uptake of bleomycin that was previously administered (locally or intravenously) at a very low dose[11]; alternatively, cisplatin can be used locally with equally good results[12]. ECT is mainly indicated for cutaneous and subcutaneous tumors of any histology. As cell membrane permeabilization is produced by a physical phenomenon known as electroporation, it affects all cells, regardless of the histology of the tumor. Recently, a meta-analysis of ECT clinical studies in human oncology showed that the overall objective response (OR) rate vary from 62.6% to 82.2% depending also on the route of administration of the drug, being either intravenous or intratumoral[13]. Great efforts are being made to extend ECT to non-cutaneous locations such as the liver[14], the brain[15] and the bones[16]. Moreover, an endoscopic electrode was developed to treat the colon[17]. The considerable experience and deeper understanding gained since ECT inception prompted the publication of new and more flexible standard operating procedures[18].

Furthermore, the immune response to tumor cells is currently one of the major areas of research in biomedical science. In the case of melanoma, the evidence that a number of tumor-associated antigens (TAA) are recognized by both CD4+ and CD8+ T lymphocytes and a family of tumor-specific antigens (MAGE) has been identified, make this disease treatable by immunotherapy. The clinical experience with melanoma immunotherapies seems promising, with increasing evidence that combined approaches may be required to ensure durable responses in a majority of patients[19] In vivo ECT treatment has proven to possess an intrinsic cytotoxic property toward cancer cells, and also to generate a systemic anticancer immune response via the activation of immunogenic cell death[20]. For these reasons, this drug delivery method can represent an important strategy to propose as a single regimen or in combined therapeutic protocols.

In this translational study, we present results obtained with 67 canine patients affected by oral malignant melanomas at different stages, treated with ECT and with a follow-up of two years. We also discuss the benefits of this treatment compared with classical therapeutic approaches, with the aim to open a perspective for human treatments at earlier stages.

## Results

Sixty-seven patients were enrolled. Differences in age or body weight were not significant among different clinical stages (p>0.1). Median age was 12 years (6-16 years) and median body weight was 22 kg (3.5-58 kg). Most common breed was crossbreed (24), followed by Labrador retriever (9), cocker spaniel (7), golden retriever (6), beagle (4), poodle (4), Rottweiler (3), Doberman (2), shar-pei (2), dogo (1), English Mastiff (1), chou chou (1), basset hound (1), Pekingese (1), and Dalmatian (1). The number of patients in stage I was 11, in stage II 19, in stage III 26, and in stage IV 11.

Among all the tumors, 52.2% had a rostral location, and 46.3% a caudal location, while 1.5% (1 case) was located in the root of the tongue. Patients with tumors in a caudal location had bone involvement more frequently (55.8% vs 44.2%, bone involvement caudal vs rostral, respectively). Also, patients with tumors in a caudal location developed new metastases more frequently (35.5% vs 22.2%, percent of patients who developed new metastases caudal vs rostral location of the tumor, respectively). Also, the local response was worse in these patients, where 58.1% OR were obtained (CR 16.1%, PR 41.9%, SD 22.6% and 19.4% PD) when compared with tumors in a rostral location (OR 80%, CR 25.7%, PR 54.3%, SD 11.4% and PD 8.6%).

The frequency of histologic subtypes: 25.4% epithelioid, 25.4% mixed, 25.4% spindle cell, 13.4% anaplastic and 10.4% others. Six cases were amelanotic melanomas.

Kaplan-Meier curves of survival were significantly different in all groups (p<0.05). See Fig. 1.

**Figure 1.**
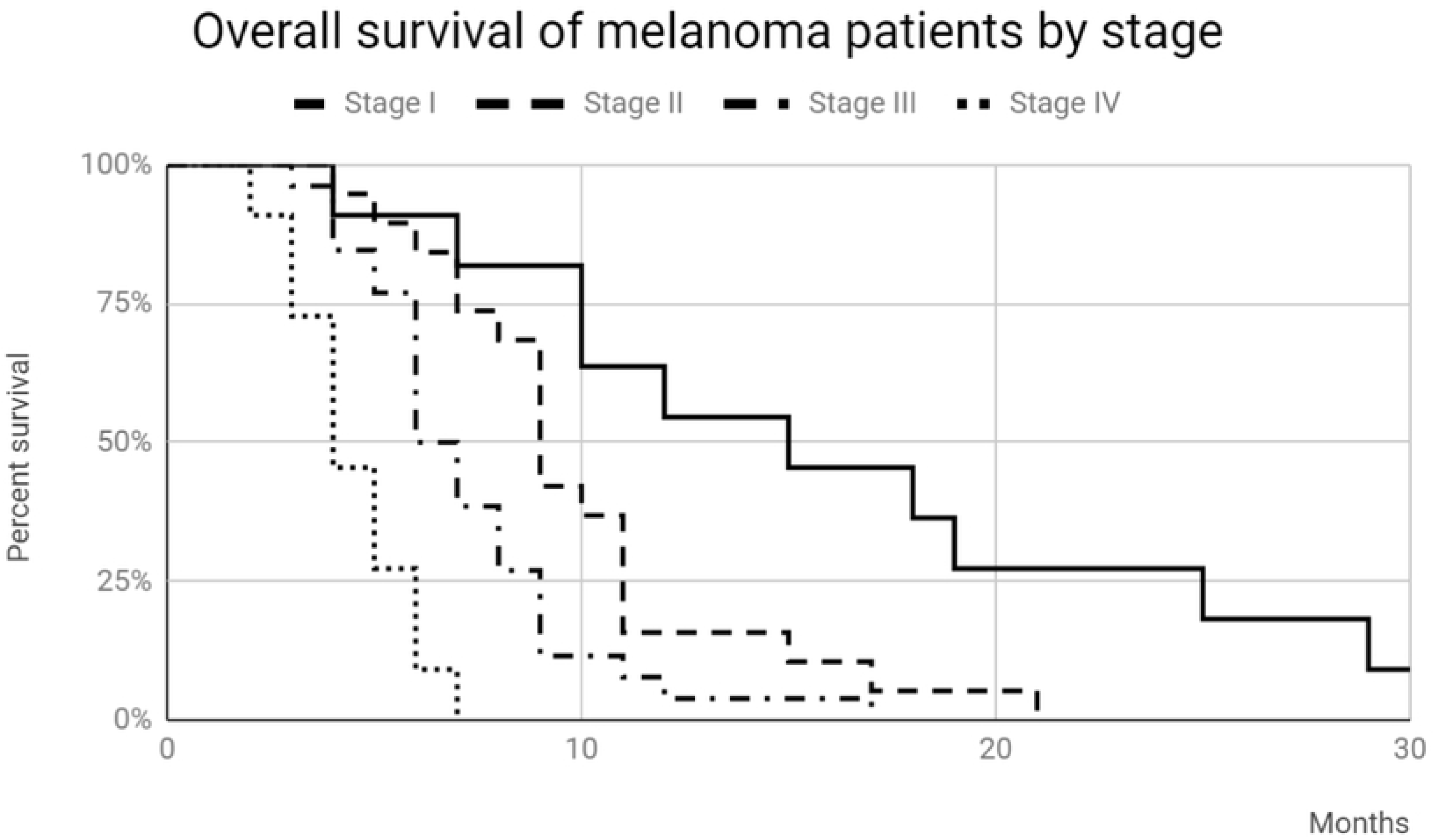
Kaplan-Meier curves of survival for the patients in each stage (p<0.05).

The overall median survival in all stages was 7.5 months (2-30 months, mean 9.1 months).

Median survival of patients in stage I was 16.5 months (4-30 months, mean 16.27 months), stage II was 9 months (4-21 months, mean 9.95 months), stage III 7.5 months (3-17 months, mean 7.23 months) and stage IV 4.5 months (2-7 months, median 4.45 months).

The average number of ECT sessions was 1.5 for every stage.

According to WHO criteria the objective overall response (OR) rate was 70.1%; with 20.9% complete responses (CR), 49.3% partial responses (PR), 16.4% stable disease (SD) and 13.4% progressive disease (PD). The local response in each stage was (see Fig. 2):

**Figure 2.**
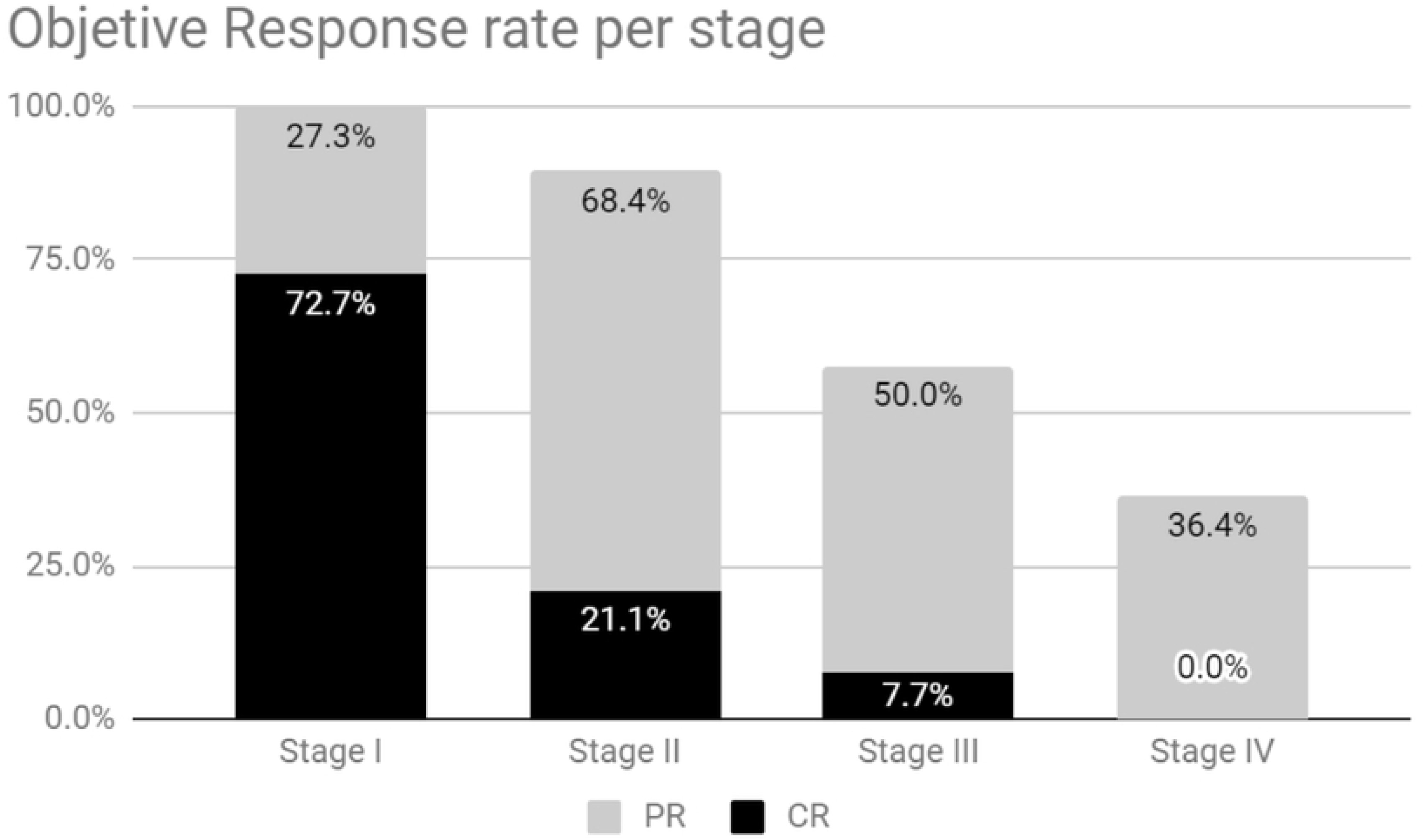
The bars show partial and complete response rate obtained in each stage.

- Stage I: CR 72.7% PR 27.3% SD 0% PD 0%, OR 100%.
- Stage II: CR 21.1% PR 68.4% SD 0% PD 10.5%, OR 89.5%.
- Stage III: CR 7.7% PR 50% SD 26.9% PD 15.4%, OR 57.7%.
- Stage IV: CR 0% PR 36.4% SD 36.4% PD 27.3%, OR 36.4%.

Bone involvement determined by radiology and/or CT scan, was present in 64.2 % of the patients. The most common histological subtype with bone involvement was anaplastic, followed by mixed and spindle cells.

Overall survival in patients with bone involvement was lower, with a median of 3 months (2-18 months, mean 5.6 months) than in patients without bone involvement, with a median of 11.5 months (4-30 months, mean 14.3 months). Kaplan-Meier curves show that the difference is statistically significant (p<0.01), see Fig. 3.

**Figure 3.**
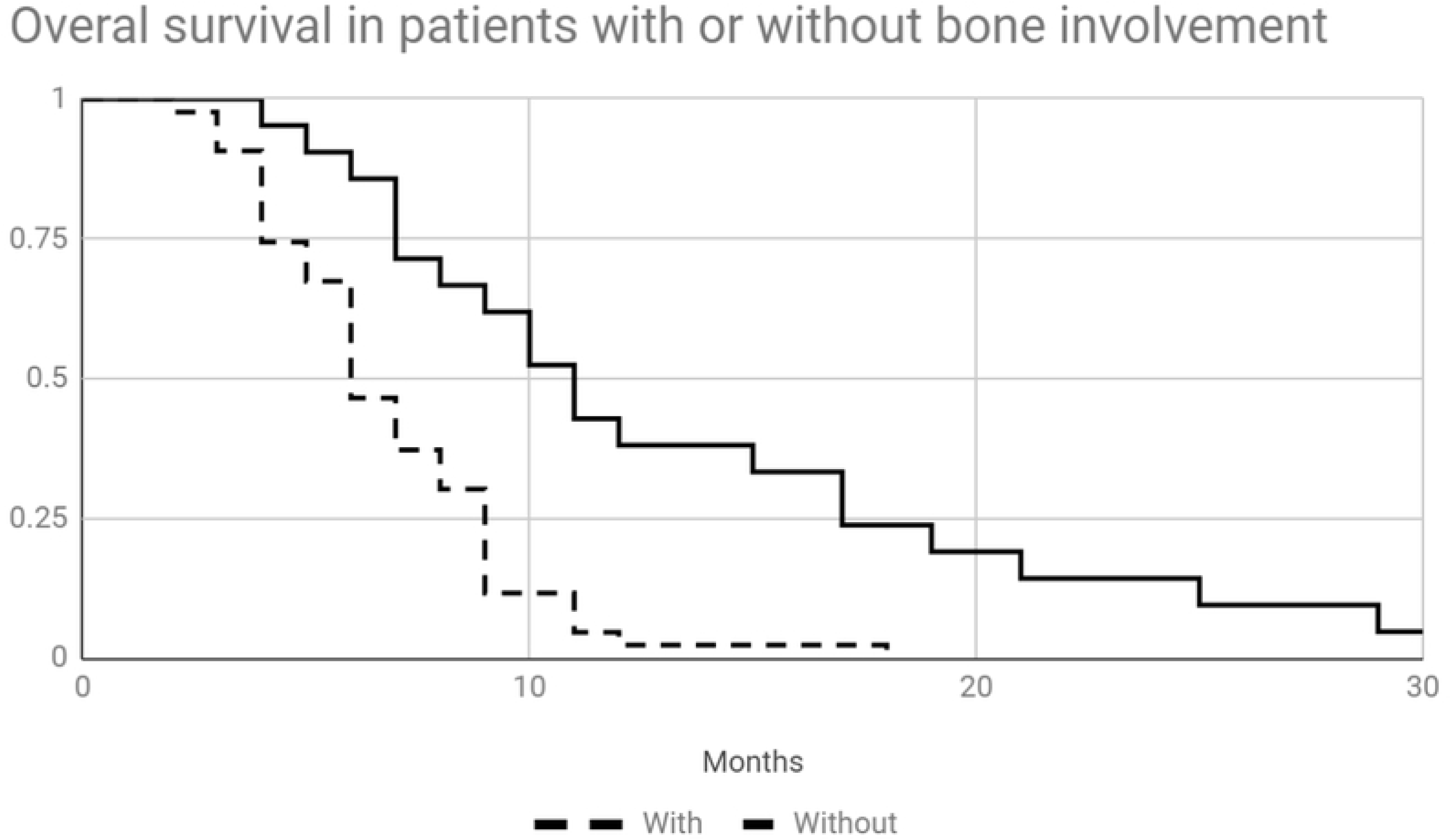
Kaplan-Meier curves of survival comparing patients with and without bone involvement (p<0.01).

Also, local response rates in patients with bone involvement were poorer with 60.5% of OR (CR 9.3%, PR 51.2%, SD 20.9% and PD 18.6%) vs 90.5% (CR 47.6%, PR 42.9%, SD 9.5% and 0% PD), see Fig. 4.

**Figure 4.**
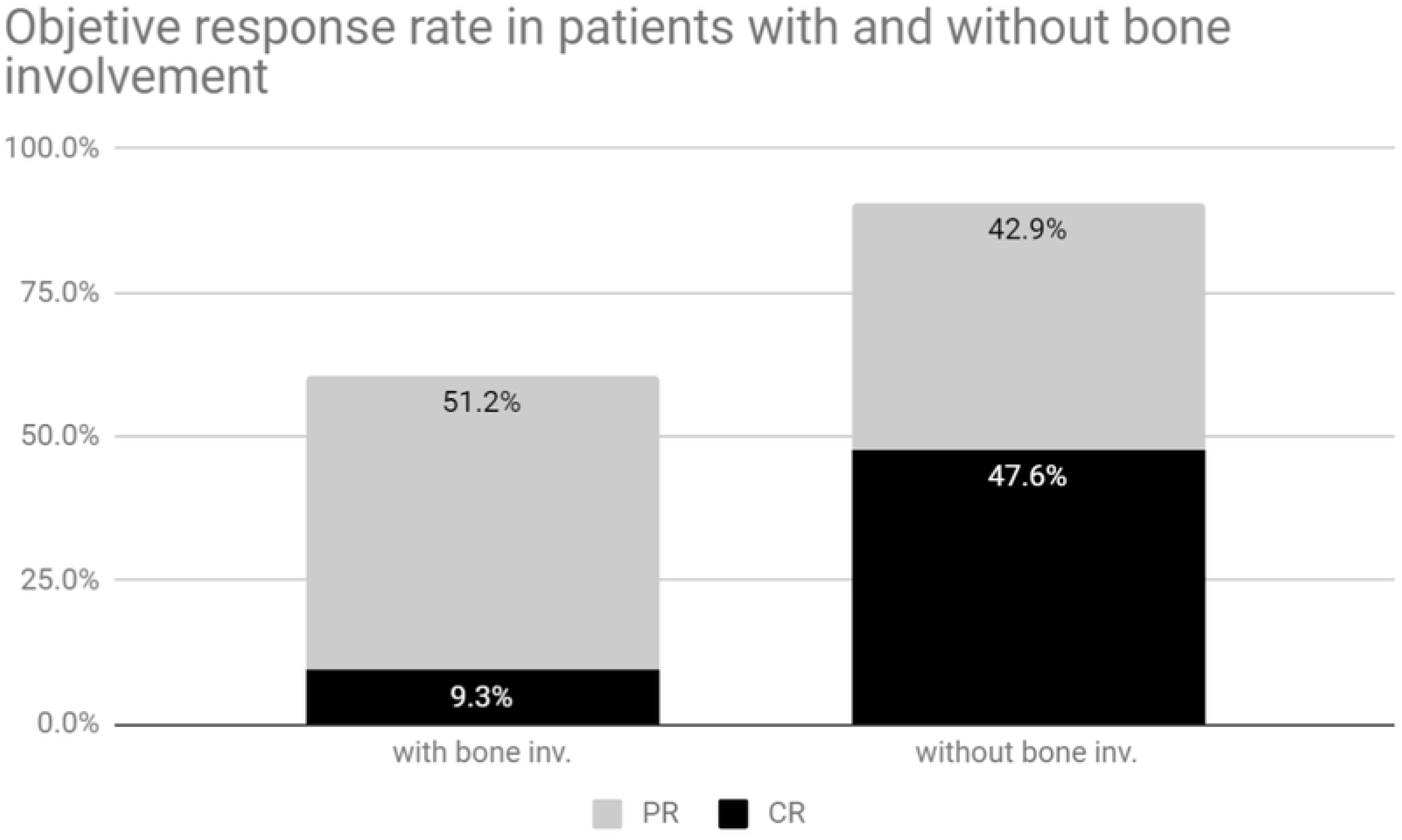
The bars show the sum of partial and complete response rates in patients with and without bone involvement.

Among the treated patients, 31.3% had metastases at presentation, 52.3% of these corresponding to lymph node metastasis. Among the patients that did not have metastasis at the time of diagnosis, 23.9% developed new metastases.

Among the patients that developed new metastases, 73.7% had bone involvement at the time of diagnosis (see Fig. 5.) and 57.9% had a caudal location (see Fig. 6). The anaplastic histological subtype was the most frequent tumor that developed new metastases, followed by spindle cell and epithelioid. Patients treated in stage I did not develop new metastases, but in the other stages, the incidence of metastases was similar, around 30% (31.6%, 36.8% and 31.6% for stages II, III and IV, respectively).

**Figure 5.**
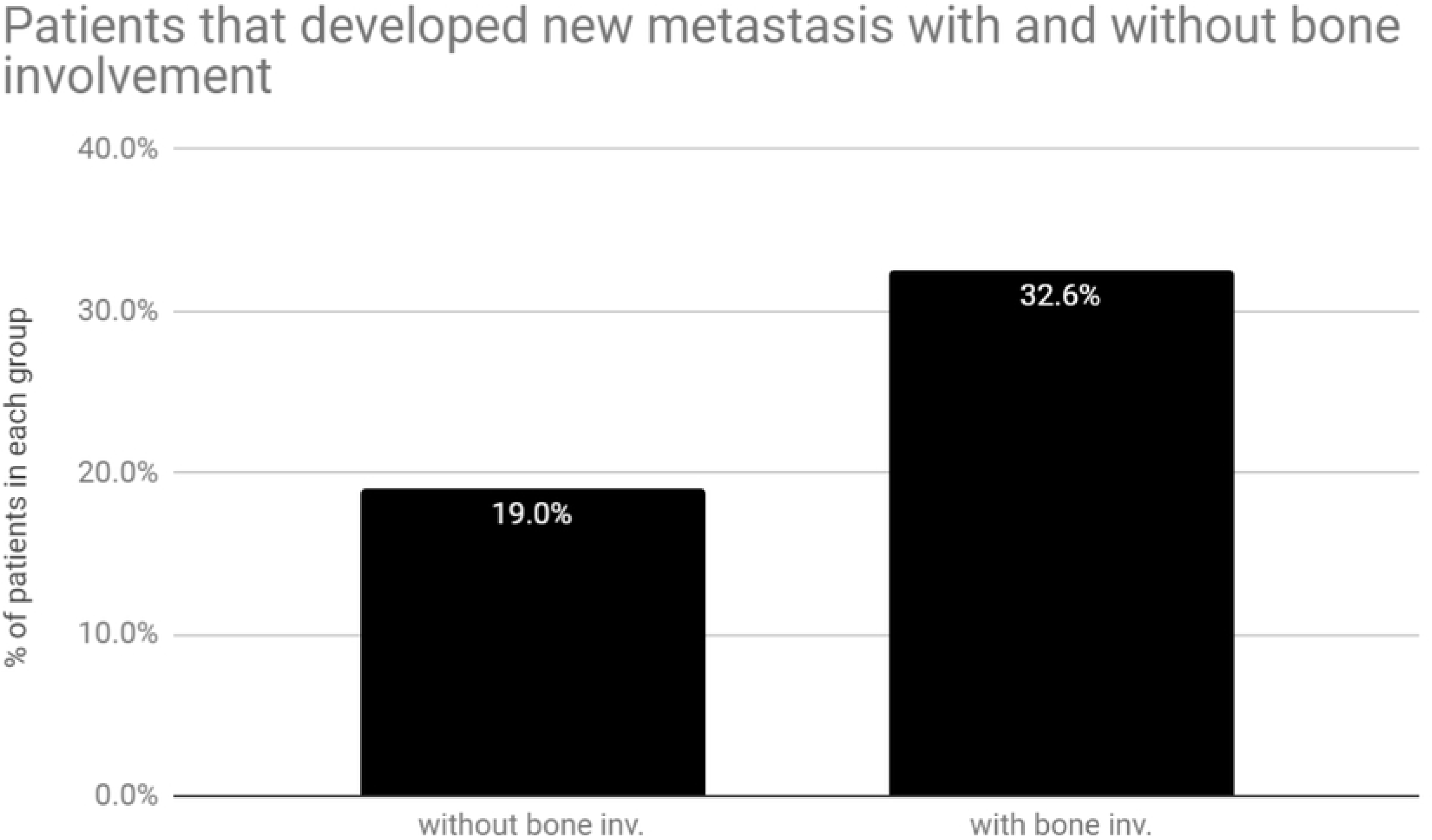
The bars show the percentage of patients that developed new metastases when there was bone involvement vs when it was not.

**Figure 6.**
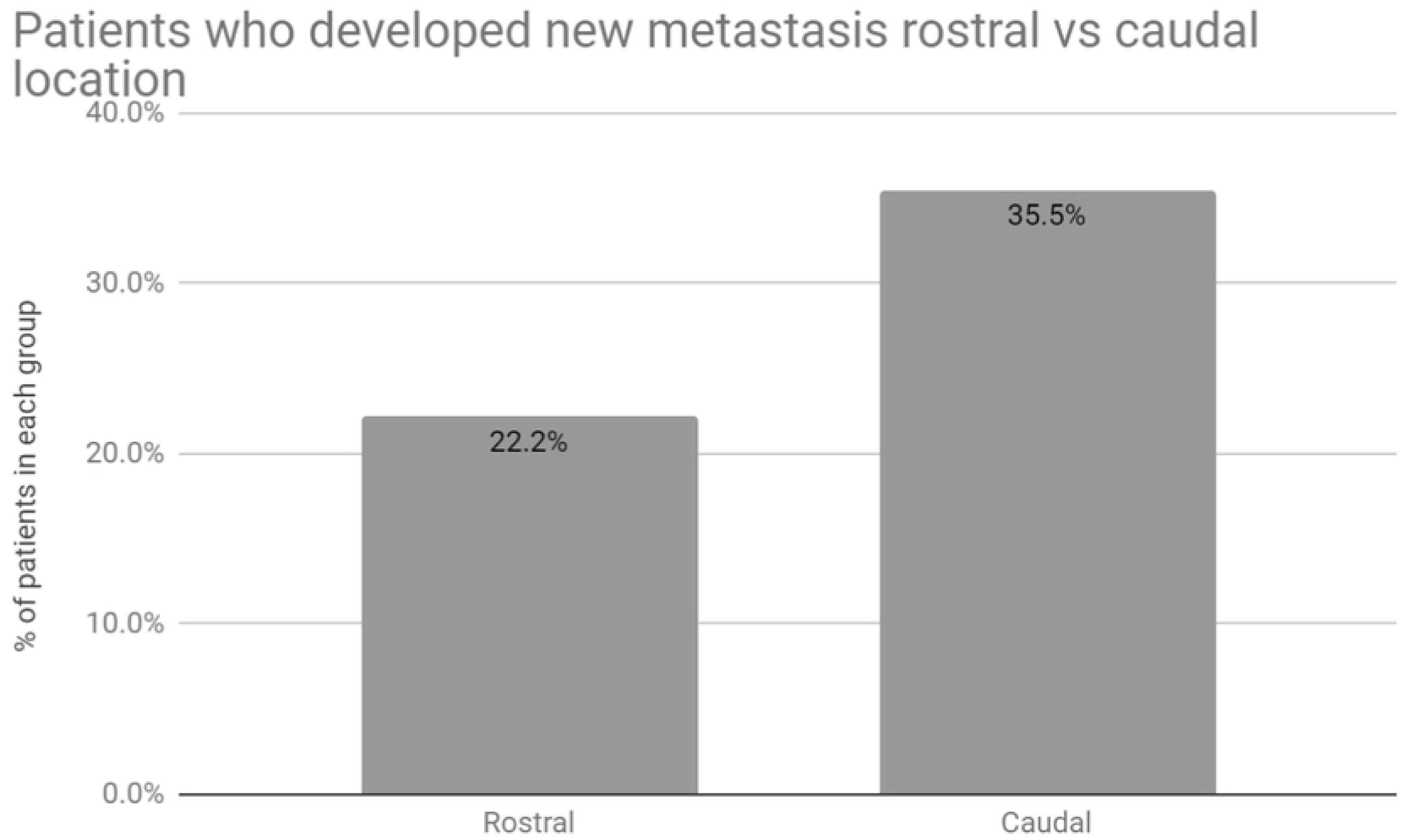
The bars compare the percentage of patients that developed new metastases with tumors in a rostral vs caudal location.

The main cause of death during this study was euthanasia accounting for 77.6% of deaths, where 49.3% corresponded to euthanasia due to local progression of the disease and 28.4% due to metastatic progression of the disease. Unrelated causes accounted for 17.9% and 4.5% of the patients that remained alive at the end of the follow-up period (24 months).

Due to the fact that the lesions were located in the oral cavity, many symptoms that affected the quality of life were present. Among them: bleeding, accidental biting of the tumor, pain, impossibility to feed adequately (see Fig. 7). Quality of life assessment revealed that, patients in stage I with a tumor in a rostral location had no negative impact in their quality of life and for that reason, even if symptoms like bad smell and bleeding ceased, there was no change in it. Patients in stages II, III and IV that obtained a partial, or complete responses did have an improvement in their quality of life, but the ones that obtained a stable or progressive disease response, did not.

**Figure 7.** Picture of the oral cavity of one of the canine patients treated, with a stage II caudal malignant melanoma. In a, before treatment, as can be seen the mass is located in the soft palate, it greatly affected the quality of life of the patient because it provoked eating difficulties, pain and bleeding. In b, a PR was obtained, although a fistula was formed, quality of life improved as eating difficulties disappeared. The fistula rendered no symptoms.

## Discussion

In this study, we describe the largest group of dogs with oral malignant melanoma treated with electrochemotherapy with a 2-year follow-up with the aim of not only providing results of a veterinarian study but of a translational study[21], as canine oral melanomas share many resemblances with human melanomas and are considered a very good preclinical model[22].

The frequency of breeds, no differences in gender predisposition and age at the diagnosis are in accordance with different authors[23][24]. In agreement with these results, most of the patients who attend our clinic in Argentina are crossbreed. (Unpublished data).

Most of the patients in this study were in stage II or III. We believe that this is because the owner was not able to recognize the lesions earlier in the onset of the disease. As the lesions are located inside the oral cavity, and first symptoms are related to changes in color, shape, characteristics of previous nevii or new lesions that are very small, owners make a visit when other and more evident symptoms of the disease appear, which are more common in advanced stages as they are related to the size of the lesion

Many patients in stages III and IV were treated with non-oncological conservative surgery and then attended our clinic with local or metastatic recurrence[25].

The most frequent rostral location of the melanoma makes it easier for the owner to detect it and seek for advice. This delay in the visit provided more time for the bone invasion to occur and makes it more prone to metastasize for the same reason.

In our study, melanomas located in the tongue are rare as reported by other authors[26].

Kaplan-Meier survival curves for comparing rostral versus caudal location revealed no statistically significant differences in accordance with other authors (data not shown)[27].

We had a similar incidence of histological subtypes to the one reported by Ramos-Vara et al., except for a lower incidence of spindle cells type. Also, we had a higher incidence of amelanotic melanomas[28].

In this study, we observed that patients with bone involvement have a poorer prognosis, this could be related to a limitation of electrochemotherapy. It is known that the electric field distribution in the bone is different than in the tissue, providing an inhomogeneous electric field. This situation can reduce the adequate permeabilization of cells, and for this reason, the bone would not be adequately treated during the ECT session. Moreover, inserting the needles in bone tissue is often very difficult and sometimes impossible, especially in early phases of bone invasion; this also explains why more objective responses were obtained in patients without bone involvement (90.5% vs 60.5%). This situation makes the adequate coverage of the tumoral tissue and margins impossible. Treating bone every time its invasion is suspected with the adequate equipment may have changed the outcome of this study. Also, we must mention that this poorer prognosis could be also due to the fact that patients with bone involvement were in more advanced stages of the disease. Other authors found better results with small, rostral tumors without bone involvement[29–31].

If we consider patients in stage II, and analyze the response with and without local involvement, we observe that patients without bone involvement had a better local response (100% OR vs 81.8% OR, without and with bone involvement respectively). Therefore, we suggest that bone involvement is an important factor to take into account for treatment selection and consider the limitations of ECT for these cases. Additional data is needed to confirm this hypothesis.

In human TNM staging for mucosal melanoma, bone or cartilage invasion is T4 regardless of the size of the lesion[32]. If we apply this criterion, all the patients with bone involvement will be staged as IV. In this work, we found that the local response and survival times of patients in stage III were similar to the patients with bone invasion, and for that reason, we consider that patients with bone invasion should be staged as III, regardless of the size of the lesion.

Overall survival of the patients varied according to the stage as it was expected. We believe that this is related not only to the disease but to the effectiveness of electrochemotherapy in each stage. As a local treatment, best results are obtained when no dissemination is present. Also, smaller lesions can be treated more easily and covered by the 6-needle electrode. Bigger lesions may present a necrotic center or may have poorly irrigated areas leading to a poor distribution of the drug in the tumor.

In relation to lymph node involvement, many patients with lymph nodes that were normal to palpation were positive in fine needle aspiration samples. In these cases, good local response was obtained with a stable systemic response. Nevertheless, patients with enlarged lymph nodes were subjected to lymphadenectomy after electrochemotherapy to improve their quality of life.

When euthanasia is possible, the survival time of the patient is highly influenced by the quality of life. In more advanced stages of the disease, there is a serious commitment to improve the quality of life of the patients; since achieving this will prolong survival.

The average number of ECT sessions, until a steady response was reached, was 1.5 (range 1 to 4), regardless of the stage. This result was not anticipated, rather we were expecting to have an average of more sessions in more advanced stages, but survival time may not allow to perform more sessions. Factors that we consider influence the number of sessions are: 1) The size of the tumor (as the adequate coverage of the tumor with the electrodes becomes more difficult in bigger tumors), in fact, the updated standard operating procedures for electrochemotherapy recommend moderate overlapping of the electrodes[18], an approach that was discouraged in the previous standard operating procedures for normal tissues. 2) In relation to vascularization, tumors growing very fast tend to have necrotic areas with poor vascularization[33], this makes the distribution of the drug more difficult. To address this problem, we recommend the use of a combination of local and systemic bleomycin in big or in very heterogeneous tumors[34]). This drug combination can be performed in any of the ECT sessions, and for that reason, patients that had disease progression before the first month following the ECT session received a combined administration of bleomycin.

Best local responses were obtained with patients in stage I and II, with 100% and 89.5% of OR. CR rates were higher in patients in stage I, and PR in patients in stage II. While in stage III, the percentage of SD response rate is higher than in previous stages. In this stage, patients with and without lymph node involvement are grouped. If we analyze local response with and without lymph node involvement, they seem to be very similar. In fact, in our work, patients with lymph node involvement had better responses. Because of this, we believe that tumor size is more important than lymph node involvement for predicting the response to electrochemotherapy, and lesions of less than 4 cm in diameter yield the best results when no metastases are present.

In our study, the incidence of metastasis was lower than in other studies. Patients in stage I did not develop new metastases probably due to local control of the disease early in its course. We observed that bone involvement was present in most of the patients that later developed new metastases. This could be due to the difficulty in treating the invaded bone, and thus this untreated area could lead to a dissemination of tumoral cells later on the progression of the disease. Also, caudal location is in general more difficult to reach and to cover it adequately with the electrodes. Finally, the anaplastic histological subtype was more correlated with the development of new metastases, but in this case, this could be due to the higher aggressiveness of this histological subtype.

As mentioned before, the quality of life of the patients is very important, and poor quality of life is an indication of euthanasia. For this reason, survival times are highly influenced by it. In this study, as a very good local response was obtained, quality of life improved and because of that, survival times too, as happens with other treatments that achieve good local control.

Surgery is the most common treatment to manage the local tumor. The median survival time for dogs with malignant melanoma treated with surgery alone varies from 5 to 11 months[35–37].

Conservative surgery is only recommended followed by radiotherapy in non-resectable tumors, for local control of the disease. Electrochemotherapy is an excellent option to perform alone or postoperatively with similar results.

For stages I to III, survival median time is 7.6 months[38].

Median survival times are similar to ours, and that was expected, since both are treatments that provide only local control of the disease. When surgery is possible and the owner accepts it, we recommend it, as surgery is a well-established technique in our setting. Electrochemotherapy can be used to extend surgical margins, or even to treat the scar to avoid a recurrence. We chose to perform electrochemotherapy as a first-line treatment only when surgery is not possible. This is because local tumor recurrence is more frequent following incomplete resection with about 62%–65% recurrence rate (15%-22% recurrence rate following complete excision)[39].

Radiation therapy is effective in obtaining good local control of the disease, especially with patients that are not suitable for complete surgical excision. The response rates are similar regardless of the protocol used. The addition of chemotherapy provides no additional benefit. Proux et al. achieved 51% CR, 31% PR, 16% SD 1% PD. They report obtaining more complete responses followed by partial responses. In our case, if we consider all stages together, more partial responses than complete ones were obtained. However, despite of this very good local control of the disease, they report 51% incidence of new metastases, while, in our study, the incidence of new metastases was 29.3%. Results of median overall survival were 7 months (6-9 months), similar to our 7.5 months (2-30 months), but the range is much wider in our case[40].

It is well known that melanoma is an extremely chemotherapy-resistant tumor, and recent studies point out that using this approach as adjuvant to other treatments has little or no benefit. Boria et al.[41] reported a tumor remission of 18% in melanoma patients and a median survival time of 4 months. In this sense, electrochemotherapy consists of a new way of delivering chemotherapeutic drugs, in particular, bleomycin, that dramatically increases the effectivity of chemotherapy. But it only works as a local treatment. Although this treatment seems to be effective only locally, lots of benefits for the patient, and consequently for the owners, have been shown, making this approach very promising in veterinary and human therapeutic applications.

It is very clear that the treatment of canine oral malignant melanoma requires an effective local treatment as the first step to improve the control of the symptomatic disease, improve the quality of life and delay euthanasia. However, it would be beneficial to perform a systemic adjuvant therapy to further improve the survival of treated patients.

A retrospective study compared survival times in 140 canine oral melanomas treated with external beam radiation with and without chemotherapy. They found no added benefit with it[42].

Because the results are not conclusive regarding chemotherapy, new immunological therapies are being evaluated. As canine malignant melanoma shares the mechanisms with human melanoma for immune system response evasion, it is a very attractive target for immunotherapy[43].

New immunological approaches are now available to improve the results of local therapy. Most of the patients have lymph node involvement or microscopic dissemination of tumoral cells that are not detectable at the time of diagnosis.

Promising results have been published by Mozillo et al. combining ECT with ipilimumab. The ECT session was performed for local control of the disease after treatment with ipilimumab. It seems that ECT triggered a systemic response, which was obtained in 60% of the patients[44].

In the last decade, DNA-based immunotherapy has reached important achievements in the treatment of infectious diseases and cancer[45].

In other work, we explored the use of ECT in combination with canine IL-2+IL-12 plasmid vector delivered by electroporation, obtaining local control of the disease in patients with different histologies. We observed fever, swelling, and lethargy due to the transfection, which had to be treated with corticosteroids. A single systemic response in lung metastasis was obtained[46]. We continue working in a protocol combining GET with ECT to enhance the immune response and thus improve treatment response and induce an abscopal effect. So far, we observed that in the oral cavity, only SD had been achieved (data not shown).

Cemazar et al. combined ECT with human IL-12 plasmid vector delivered by electroporation and obtained a 72% CR rate in spontaneous canine mast cell tumors without side effects. They also observed the treatment prevented local recurrences and distant metastases[47].

The possibility of using an IL-12 plasmid without an antibiotic resistance gene was explored by Tratar et al.[48] in a canine melanoma model. They found that this plasmid has similar cytotoxicity and expression profiles to the plasmids that encode the antibiotic resistance gene. Also, by using constitutive-specific promoters, they obtained better results than using fibroblast specific promoters.

Bergman et al. showed very promising results in a small cohort of dogs receiving DNA Vaccination with Xenogeneic Human Tyrosinase[49]. This led to the development of Oncept™ vaccine, with a reported overall median survival time of 19 months for patients in stage II/III[50]. But after the FDA approval, opposite results were obtained with the use of this vaccine, as Grosenbaugh et al. did not reach the specific median survival time[51], and Ottnod et al. found no differences in progression free survival and overall survival rates between vaccinated and non-vaccinated patients[52]. A randomized, prospective, double-blinded clinical trial is needed to define whether or not the use of Oncept™ is recommended[53].

The use of animal models to predict human response to therapies has become one of the main focal points of the scientific community. Pet animals, especially dogs, represent an excellent model for anti-cancer treatments. In fact, human and dogs’ tumors present many similarities: they develop spontaneously in older subjects and are histologically comparable; they share conventional treatments with similar responses. Differently than induced murine models, the development of canine tumors can be observed over a long time and the life span of canines is proportionally more comparable to humans than the lifespan of rodents[54,55]. Because melanomas in dogs and humans show very similar characteristics, we present these results to the scientific community, with the aim to achieve soon successful translational therapeutic protocols by considering the use of electrochemotherapy.

### Patients and Methods

In the present study, 67 canine patients were enrolled in this ECT protocol for oral mucosal melanoma. These patients had little chances to perform complete surgical resection due to the impossibility of clean margin resection, or because their owners rejected radical surgery. In all the patients we performed a complete physical examination, complete blood count, and serum biochemical analysis. The size of the tumor was measured using a caliper or calculated by CT scan. Dogs were evaluated for metastatic disease via 3-view thoracic radiography, abdominal ultrasound and cytologic examination of fine-needle aspirate samples or histopathological evaluation of regional lymph nodes. CT scan was performed in cases which the tumor involved deep structures such as the retro orbital region, the nasopharynx or the nasal cavity, and to confirm bone involvement.

Among the patients, 11 were in stage I, 19 in stage II, 26 in stage III and 11 in stage IV. The age of the patients ranged from 6 to 16 months, with an average of 11.7 months. There were 37 males and 30 females. The most common breeds were: cross-breed, Labrador retriever, cocker and golden retriever.

The response to the treatment was evaluated according to WHO criteria.

All the regulations from the Consejo Profesional de Medicos Veterinarios were followed. Informed consents were signed by the owners. The protocol for the treatment was approved by the CICUAL of the Faculty of Veterinary Sciences, University of Buenos Aires, Argentina. Protocol number: 2018/31.

Recommendations to report results in electrochemotherapy studies from Campana et al.[56] were followed.

The treatment was performed under general anesthesia with premedication with intramuscular administration of xylazine (Xylazine 100®, Richmond, Buenos Aires, Argentina) 0.5 mg/kg and tramadol (Tramadol®, John Martin, Buenos Aires, Argentina) 2 mg/kg. The induction was made with intravenous administration of propofol (Propofol Gemepe®, Gemepe, Buenos Aires, Argentina) 3 mg/kg. Maintenance was performed with inhaled isoflurane (Zuflax®, Richmond, Buenos Aires, Argentina) 2–3% and intravenous fentanyl (Fentanyl Gemepe®, Gemepe, Buenos Aires, Argentina) 2 mcg/kg. This anesthesia scheme provided adequate comfort during the treatment. Prophylactic antibiotics and meloxicam (Meloxivet®, John Martin, Buenos Aires, Argentina) 0.2 mg/kg were administered orally for analgesia after the treatment according to the needs of each patient.

The ECT procedure consisted of an intravenous bolus of bleomycin (Blocamicina®, Gador, Buenos Aires, Argentina) at a dose of 15 000 IU/m2 of body surface area in 30–45 seconds, followed by the delivery of the electric pulses eight minutes later to allow drug distribution according to the Standard Operating Procedures[57].

The patients were treated using a 6-needle electrode (two rows of 3 needles separated 8 mm from each other). Eight square pulses of 1000 V/cm at 10 Hz were applied until the whole surface of the tumor was covered. For pulse delivery, a BTX ECM 830 (Harvard Apparatus, Holliston, MA, USA) was used. The electrical current was monitored during the procedure using an Agilent DSOX2012A oscilloscope (Agilent Technologies, Santa Clara, CA, USA), confirming the adequate delivery of the pulses.

In cases of probable or certain nasal duct invasion we used the Single Needle Electrode®. It is an electrode specifically designed in our lab for deep-seated tumor treatment with ECT. It consists of an insulated needle with an anode on one side and a cathode on the other.[58]

The patients were scheduled for control at 30, 60 and 90 days, and every 3-4 months for two years, when a complete clinical examination was performed to determine lymph node enlargement. Thoracic x-rays were performed to determine the development of lung metastases or when symptoms appeared. As lymph nodes and lungs are the most common sites of metastatic dissemination[59].

Quality of life assessment was performed using a modified questionnaire from Lynch et al. [60], and following their recommendations. The data obtained by the treating oncologist was included in the medical record of the patient.

Statistical analysis was performed using Mann Whitney U test to compare the participants age and body weight.

Kaplan-Meier survival curves and Log-rank significance test were performed in order to study differences among different groups.

## Acknowledgements

F.H.M., S.D.M. and G.R.M. are researchers from Consejo Nacional de Investigaciones Científicas y Técnicas (CONICET). This work is supported by grants from CONICET, www.conicet.gov.ar, (PIP 379/2012/2016 and STAN 534/12), CNR (CNR Project DSB.AD007.072) and Universidad de Buenos Aires, www.uba.ar, (UBACyT 2014/2017). The funders had no role in study design, data collection and analysis, decision to publish, or preparation of the manuscript.

This manuscript was proofread by PurpleGY.com.

## Author contributions statement

M.N.T. and F.H.M. worked in the design of the study, in the selection and treatment of the patients, in the analysis of the data and in the writing of the manuscript. S.D.M. worked in the treatment of the patients, in the analysis of the data and in the writing of the manuscript. G.R.M. worked in the writing of the manuscript, in the direction of the group and in the administration of the funds. E.S. worked in the design of the study, in the writing of the manuscript, in the coordination of the group, and in the administration of the funds. All authors reviewed the manuscript.

## Additional information

### Ethics approval and consent to participate

All the regulations from the Consejo Profesional de Médicos Veterinarios were followed. Informed consents were signed by the owners. The protocol for the treatment was approved by the CICUAL of the Faculty of Veterinary Sciences, University of Buenos Aires, Argentina. Protocol number: 2018/31.

### Availability of data

The datasets used and/or analysed during the current study are available from the corresponding author on reasonable request.

